# The entry of pinewood nematode is linked to programmed tracheal development of vector beetles

**DOI:** 10.1101/2022.10.29.514345

**Authors:** Xuan Tang, Tuuli-Marjaana Koski, Jianghua Sun

## Abstract

The transmission of some pathogens by insect vectors often requires fine-tuned biological synchronization and communication between the two parties. However, the role of tissue development in mediating this pathogen-vector coordination remains elusive. Here we investigated the association between tracheal development of *Monochamus alternatus* beetle and a notorious plant parasitic pine wood nematode *Bursaphelenchus xylophilus*, which enter and inhabit inside the tracheal system after adult beetle eclosion and then were transmitted to pine trees (*Pinus* spp.) by the beetle. We found that the tracheal systems of newly emerged adult beetles underwent morphological changes during the adult sclerotization, characterized by a remarkable increase of diameter of thoracic tracheal dorsal tubes. Consistently, comparative transcriptomics further revealed dramatic changes in gene expression occurring within five days after eclosion, demonstrating sequential regulation of tracheal genes. Genes controlling primary branching were up-regulated soon after eclosion, whereas those controlling terminal branching and branch fusion were up-regulaed at the later stage. Interestingly, during tracheal maturation, genes involved in biosynthesis of Juvenile Hormone (JH) and ecdysteroids were activated. Nematode loading assay further revealed that the entry of nematodes to beetle tracheae was initiated earliest after three days post eclosion, implying that nematode entry is reliant on the formation of the vector’s primary tracheal branches and is perhaps stimulated by production of insect hormones or their precursors. In addition, specific regulation on secreted proteins, such as Defensin-1-like and membrane proteins, at later stages of tracheal maturation may further facilitate the entry of nematodes by improving their retention. Therefore, this study highlights vital role of programed tracheal development on nematode entry into its vector beetle, and suggests a potential roles of insect genes and metabolites in manipulation of interspecific biological developmental synchronization between parasites and their insect vectors.

## Introduction

The pine wood nematode (PWN), *Bursaphelenchus xylophilus*, the causal agent of pine wilt disease, poses a global threat to pine forests due to its ability to rapidly kill healthy pine (*Pinus* spp.) trees (Mamiya, 1983; Tyler et al., 2011). Because successful transmission of PWN to new host trees relies on pine sawyer beetles *Monochamus spp*. (*M. alternatus* being the primary host in China), the synchronization of life cycles of PWN and its vector beetle has been recognized as one of crucial factors for the spread of pine wilt disease (Kobayashi et al., 1984; Togashi, 1985). Vector beetles prefer to oviposit on weakened trees (generally during early autumn) (Aikawa, 2008; Linit, 1988), and are therefore attracted to trees infested with PWN adults and larvae (L_1_–L_4_) (Akbulut and Stamps, 2012; Mamiya, 1975). If conditions inside the host tree worsen (generally during winter), L_2_ larvae molts to dispersal mode and become third-stage dispersal juveniles (L_III_) (Futai, 2013), which are attracted to the beetle larvae and aggregate around their pupal chambers (Necibi and Linit, 1998). In the following summer, L_III_ develops to fourth-stage dispersal juvenile (L_IV_) when the vector beetles metamorphose to adults inside their pupal chambers (Warren and Linit, 1993), after which the L_IV_ nematodes enter the spiracles of the beetles to reach their tracheal systems (Aikawa, 2008). This nematode loading phase lasts for several days until the adult beetles emerge and fly to feed on healthy pine trees. Once reaching the new host trees, L_IV_ juveniles invade the tree through the beetle’s feeding wounds, molt to adults, mate, and start a new PWN life cycle (Linit, 1988; Zhao et al., 2014). Understanding of the mechanisms involved in coordinating the life cycles of the nematode and the vector beetle is important for developing novel management strategies to prevent the transmission of PWN and thus pine wilt disease.

Several chemical signals have been demonstrated to be involved in the interspecific developmental cross-talk between PWN and its vector beetle. So far, it has been shown that terpene signals released by beetle larva promotes aggregation of L_III_ around their pupal chambers (Zhao et al., 2007), whereas ascarosides produced by the aggregated L_III_ promote beetle pupation (Zhao et al., 2016). In addition, fatty acid ethyl esters released from the surfaces of late staged pupae and newly eclosed adult beetles induce the molting of pupal chamber inhabiting L_III_ dispersal stage to the tracheal entering L_IV_ dispersal stage (Zhao et al., 2013). Despite that the entry of L_IV_ to beetle tracheal system is pivotal for the synchronized life cycles of PWN and its vector, the signals triggering the entry of L_IV_ to tracheae have not been investigated. The newly emerged adult beetles require the one-week period for sclerotization (Linit, 1988), and after this process, the adult beetle gnaws a gallery from the pupal chamber to the bark surface to exit the damaged tree. Thus, the tracheal loading of the newly formed L_IV_ occurs during beetle sclerotization and the process of tracheal development during this period may provide cues for PWN loading.

Insect tracheal system is an epithelial network of branched tubules extending throughout the body and the tracheal development during embryogenesis, larval and pupal stage has been intensively investigated in *Drosophila* (Ghabrial et al., 2011; Schottenfeld et al., 2010; Whitten, 1972). Tracheal development begins with the invagination of tracheal placodes into the body to form the tracheal pit during the embryonic stage (Kondo and Hayashi, 2013). The invagination phase is followed by a stereotyped primary branching that is mainly regulated by the expression of *trachealess* (*trh*) and *breathless* (*btl*) genes, which respectively encodes the bHLH-PAS transcription factor and the fibroblast growth factor (FGF) receptor (Isaac and Andrew, 1996; Wilk et al., 1996). The primary branches migrate towards their corresponding target tissues led by tip cells and successively sprout secondary branches (Affolter and Shilo, 2000). Then the tip cell specifies a fusion cell fate controlled by combined action of Bnl/FGF and Wingless signaling, increasing the expression of *escargot* (*egg*), *dysfusion* (*dysf*) genes (Jiang and Crews, 2003; Samakovlis et al., 1996). At the same time, the neighboring cell acquires terminal cell fate by expression of ERK and *blistered* gene (Best, 2019). The fusion cells migrate to contralateral sides and establish adhesion to each other to form interconnected tracheal network (Caviglia and Luschnig, 2014). In the larval stage, the terminal cells send out cytoplasmic branches (tracheoles), which cover and attach to the target tissues, a process which is known to be controlled by a hypoxia-inducible factor-α homolog, called Sima (Centanin et al., 2008). During both embryo and larval stages, the tracheal tubes undergo dramatic dilation and elongation until they reach their final functional diameter and length to fit the growth body. Apical extracellular matrix (aECM) and septate junction components have been shown play crucial role in this process (Öztürk -Çolak et al., 2016). During metamorphosis, the tracheal system is also extensively remodeled by imaginal cells as well as some multipotent larval tracheal cells (Weaver and Krasnow, 2008). Although tracheal patterns during pupal and adult stages have been described in previous studies in other insects, such as *Tenebrio molitor* (Rao et al., 2015; Ras et al., 2018), the detailed morphological and molecular mechanisms underlying tracheal development during the sclerotization process are not known.

In this study, we observed morphological changes of tracheal system in the vector beetle during metamorphosis from pupae to adult and within seven days after adult eclosion. Comparative transcriptomics of thoracic tubes at different time points post eclosion further demonstrated that tracheal systems underwent programed developmental changes at molecular level. Moreover, the Time-Series Expression Miner (STEM) analysis suggested that up-regulation of enzymes in JH and ecdysteroids biosynthesis pathways are likely central for the tracheal maturation, starting after three days post-eclosion. Consistently, in PWN loading assay, nematodes begin to enter the tracheal system at the same time point. Finally, based on the expression dynamics of differentially expressed genes (DEGs), we identified secreted and membrane proteins that potentially facilitate PWN retention after entry. These results demonstrate the key roles of tracheal maturation and related chemical cues for the entry of PWN into their vector beetles.

## Results

### Vector beetle tracheal system undergoes morphological changes after adult eclosion

To investigate the development of tracheal system in the vector beetle, we observed and compared the tracheal morphology of vector beetles during the pupal stage and the adult stage. We divided the pupal stage into four time points according to the eye color of the beetles: pupae with transparent eyes were considered to be in early pupal phase (EP), pupae with brown eyes in middle pupal stage (MP), whereas pupae with black eyes were considered to be in the beginning (LPB) or ending phase of the late pupal stage (LPE) (**Figure 1A**). Although no distinct tracheal morphological differences between EP and LPB stages were found in the microscopy images, the ambiguous tubular structures embedded in fatty tissues during LPB stage suggested that the remodeling of tracheal system occurs during this stage, and the tracheal network was formed at LPE (**Figure 1A**). To investigate tracheal development on adult beetles, we observed the tracheal morphology and measured the diameter of thoracic tracheal tubes at four time points after they emerged from pupal chambers, i.e. one, three, five and seven days post eclosion (PAE 1, PAE 3, PAE 5 and PAE 7), respectively. Microscopy investigation revealed no visible difference in tracheal morphology between beginning phase of late pupal stage and newly eclosed adults (LPE and PAE 1, respectively) (**Figure 1B**). However, significant tracheal tubule dilation was observed during the seven days post eclosion, demonstrated as a logarithmic increase in tracheal diameter (**Figure 1C**). The diameter of tracheal tubes increased nearly 30% within the first three days, but not significant increase was observed between PAE 3 and PAE 5 or between PAE 5 and PAE 7. These results suggest that most the growth of the tracheal tubes occur during the first three days post eclosion, and that reached to their final size mostly within the first five days after the adult eclosion.

**Figure 1.**
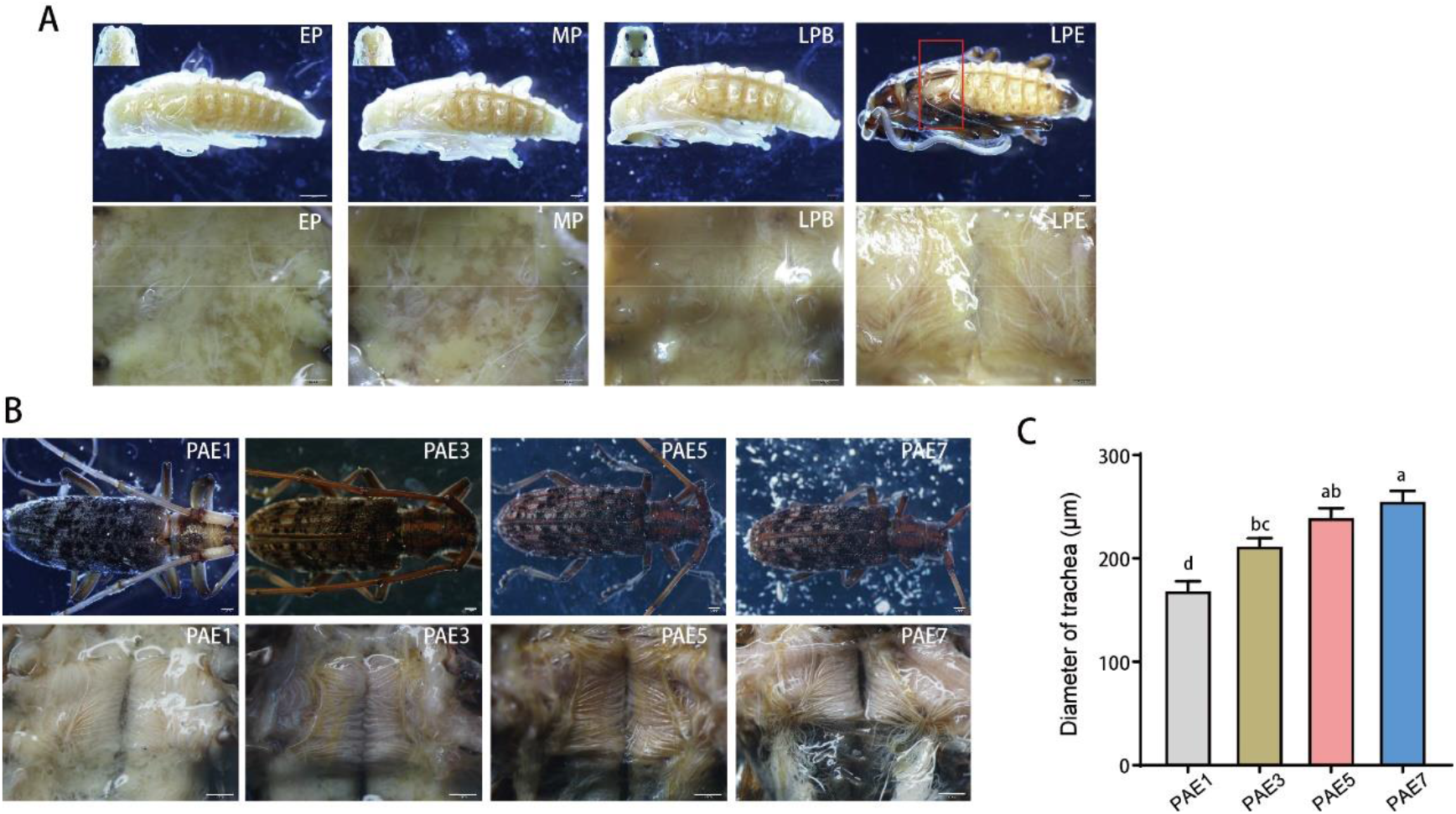
Morphological remodeling and maturation of thoracic tracheal system at different pupal stages and time points after eclosion adults in *M. alternatus*. (**A**) Representative stereomicroscopic images from four pupal stages (early stage = EP, mid-stage = MP, the beginning and end of late pupal stage LPB and LPE, respectively). Upper row present the dorsal external appearance, and lower row represent corresponding internal thoracic tracheal tubes (red frame showing the external location) (white scale bar = 2mm, black scale bar = 1mm). (**B**) Representative stereomicroscopic images from adult beetles at four different post eclosion time points (day 1= PAE 1, day 3= PAE 3, day 5 = PAE 5 and day 7 = PAE 7, respectively) for dorsal external appearance (upper row) and internal thoracic tracheal tubes (lower row, scale bar = 2mm). (**C**) Diametric expansion of thoracic tracheal tubes at PAE 1, PAE 3, PAE 5 and PAE 7. Different letters indicate statistical differences (*P* < 0.05) based on one-way ANOVA with Tukey’s multiple comparisons (*N* = 23-29 tracheal tubes from three beetles for each time point)

### Temporal changes in gene expression during tracheal development after beetle eclosion

To investigate tracheal development at molecular level, we carried out comparative transcriptome analysis of the thoracic tracheal tubes of vector beetles at PAE 1, PAE 3, PAE 5 and PAE 7. A total of 88.14 gigabytes of Illumina HiSeq data were generated from 12 samples and 69241 unigenes were obtained after assembly with an N50 value of 1438bp. The principal component analysis (PCA) based on gene expression revealed a gradual change along the principal component 1 (PC1) axis on the PC1 vs. PC2 ordination plot from PAE 1 to PAE 7 (**Figure 2A**). Samples of PAE 1 and PAE 3 were clearly separated from those of PAE 5 and PAE 7 based on PC1 score (**Figure 2A**). The number of identified genes increased from 14458 to 28441 as the trachea matured (**Figure 2B)**. Out of these genes, 6,005 and 10,632 genes were specifically expressed at PAE 5 and PAE 7, respectively (**Figure 2B)**. Differentially expressed genes (DEGs) were identified through pairwise comparison among beetles from each post eclosion time point. The number of DEGs was increased together with the difference in length of post eclosion time, i.e. the number of DEGs was highest when PAE 1 and PAE 7 were compared, and lowest when PAE 5 and PAE 7 were compared, the latter comparison resulting in only 110 up-regulaed genes with no difference in down-regulated genes (**Figure 2C**). This result is consistent with the PCA results showing no separation in expression profiles of tracheal genes at PAE 5 and PAE 7 (**Figure 2A**).

**Figure 2.**
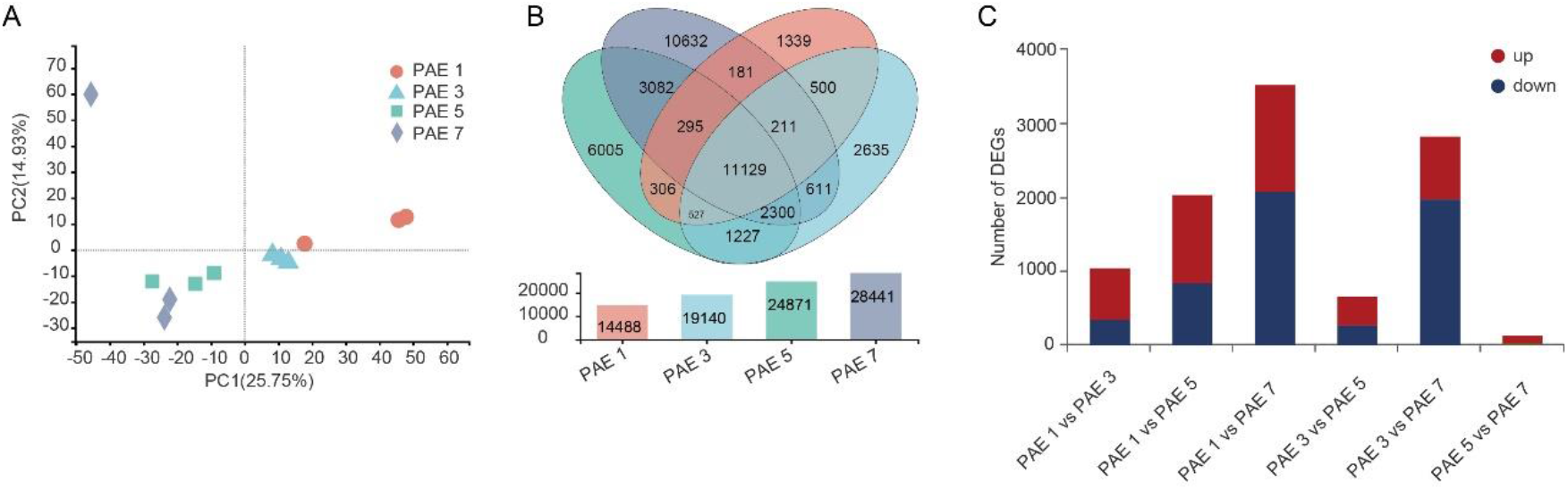
Comparative transcriptome analysis of thoracic tracheal tubes at four time points after eclosion of *M. alternatus*. (**A**) PCA analysis of thoracic tracheal tubes based on the gene expression of thoracic tracheal tubes at PAE 1, PAE 3, PAE 5 and PAE 7. (**B**) Venn diagram of identified genes expressed in thoracic tracheal tubes at PAE 1, PAE 3, PAE 5 and PAE 7. The number of identified genes in different post eclosion time points are shown below as bar plot. (**C**) Pairwise comparisons of the number of DEGs in tracheal tubes for different post eclosion time points.

### Programmed tracheal development and maturation occur within five days after adult eclosion

Based on sequence similarity to *Dropsophila* molecular marker genes controlling tracheal development (Hayashi and Kondo, 2018), as well as nr, pfam and/or KOG annotation, we identified 55 genes related to tracheal development in *M. alternatus* (**Figure 3**, Supplementary table 1). These genes were clustered based on their FPKM value across the four investigated post eclosion time points. Cluster 1 (yellow branches) included only five genes which maintained consistent expression levels between PAE 1 and PAE 5, but were down-regulated at PAE 7 (**Figure 3**). Similarly in Cluster 2 (red branches), the nineteen detected genes were up-regulated at PAE 1, but expressed at significantly lower levels after this period. These genes included *trachealess* (*trh*), a central regulator of tracheal cell fate, and *ventral veinless* (*vvl*), an additional transcriptional factor necessary for cell identity. The genes encoding FGF ligands Branchless (Bnl), FGF receptor Breathless (Btl), and its adaptor protein Dof (Stumps), were also specifically up-regulaed at PAE 1, indicating activation of Bnl-Btl signaling stimulating the tracheal branching during early stage of post eclosion. Consistently, Cluster 2 also included genes responsible for the identities of primary branch (*knirl* and *salm*) and for the migration of visceral branches (*if* and *adamTS-A*). In addition to genes involved in primary branching and its guidance, gene up-regulation was also detected in several aECM components such as Serpentine (Serp), Vermiform (Verm), and Matrix metalloproteinase 1 (MMP1) shortly after eclosion. In contrast to Cluster 2, the genes in Cluster 3 (blue branches) and Cluster 4 (green branches) were down-regulated right after eclosion (PAE 1). The 16 genes forming Cluster 3 were mainly down-regulated at PAE 1 and PAE 3, but up-regulated at PAE 5 and PAE 7. Similar to this, the 16 genes forming in Cluster 4 were down-regulated right after eclosion (PAE1), but expressed at higher levels at PAE 3 and PAE 5 (Figure 3). Both Clusters 3 and 4 included the EGF receptor (Egfr), the EGF ligand activator molecule Rhomboid (RhoA), and the intracellular signal transducers ERK (Dsor1), indicating EGF signaling is activated on day three or five after eclosion. We also discovered two genes, *hedgehog* (*hh*) and *decapentaplegic* (*dpp*) in Cluster 3, which are known to restrict the extent of migration of the dorsal and terminal branches. Notably, up-regulation of genes encoding key regulators of terminal branching (*wingless, ikke, btsz* and *silk*), branch fusion (*aop, esg, stac, CG13196*) and gas generation (*uif*) was observed after three or five days post eclosion in Clusters 3 and 4. In addition, some of the genes in Cluster 3 and 4 are likely associated with tube geometry, such those related to axial elongation (*yokie, aPKC, rhoA, shrb, yrt* and *scrib*) or encoding aECM components (*dumpy* and *mmp2*). In conclusion, despite the complexity of expression dynamics of tracheal genes during tracheal maturation post eclosion, the expression of most genes changed dramatically in an organized manner within the first five days after the emergence of the adult vector beetle. This result is also consistent with a very low number of DEGs identified in the pairwise gene expression comparison in thoracic tracheal tubes of beetles between five and seven days post eclosion (PAE 5 and PAE7, Figure 2C).

**Figure 3.**
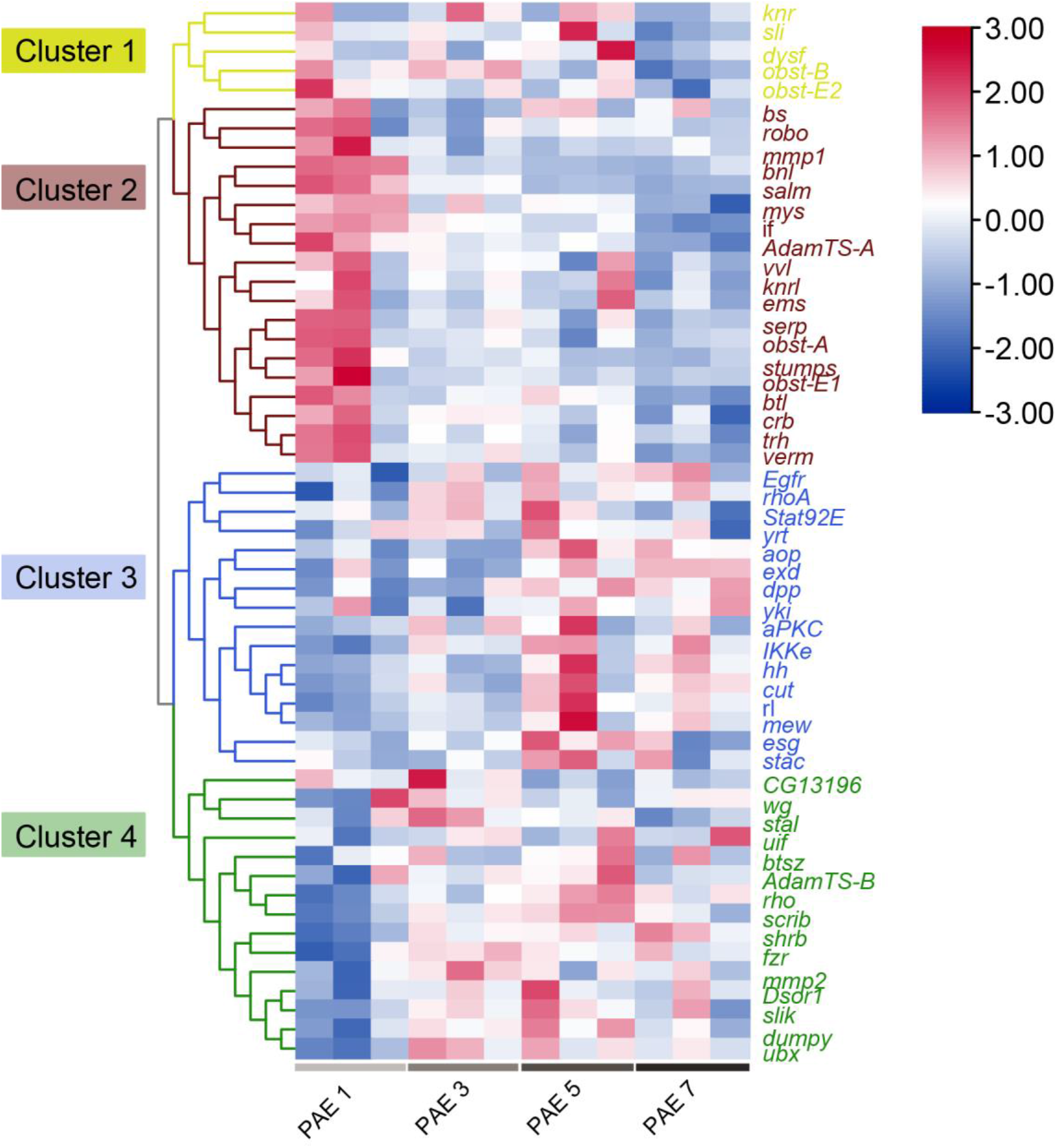
Heat map of tracheal gene expression in tracheal tubes of *M. alternatus* at PAE 1, PAE 3, PAE 5 and PAE 7. Each column represents tissues from a single biological replica (beetle individual). Differently coloured branches represent gene clustering based on the similarity of their FPKM values. Red colour indicates up-regulation and blue down-regulation of gene expression. Gene names are shown on the right.

### Hormone biosynthesis and organic acid metabolism were substantially activated in tracheal system after adult eclosion

We further identified genes which expression pattern changed most drastically during post eclosion, i.e. during tracheal maturation. The Time-Series Expression Miner (STEM) recognized nine significantly different patterns of gene expression. The expression changes in patterns 21, 24, and 25 was the most significant, and were classified into Profile 1, applying to 8914 genes. These genes were up-regulated after three, five or seven days post eclosion and maintained high until the end of the experiment (**Figure 4A**). Their Gene Ontology (GO) terms belonged to organic acid-associated metabolism, including fatty acid oxidation, and carboxylic or organic acid catabolic process (**Figure 4B**). Consistently, the KEGG enrichment analysis from these genes identified pathways related to fatty acid metabolism. Interestingly, insect hormone biosynthesis was the most significantly enriched KEGG pathway (**Figure 4C**, Supplementary table 2). The 33 genes enriched in insect hormone biosynthesis pathway were related to biosynthesis of Juvenile hormone (JH) and ecdysteroids, out of which 29 genes produce enzymes that catalyze the four key steps in JH biosynthesis (**Figure 4D**). However, in tracheal RNA libraries, we found traces of transcripts encoding JHAMT and JH epoxidase, two key enzymes in the production of JH III (**Figure 4D**). Therefore, these results imply substantial production of JH precursors during tracheal maturation after eclosion. Moreover, the expression of four genes encoding key enzymes on ecdysteroids (20-hydroxyecdysone, 20-E) biosynthesis exhibited co-varied temporal changes during maturation and were significantly up-regulated soon after eclosion (**Figure 4D**).

**Figure 4.**
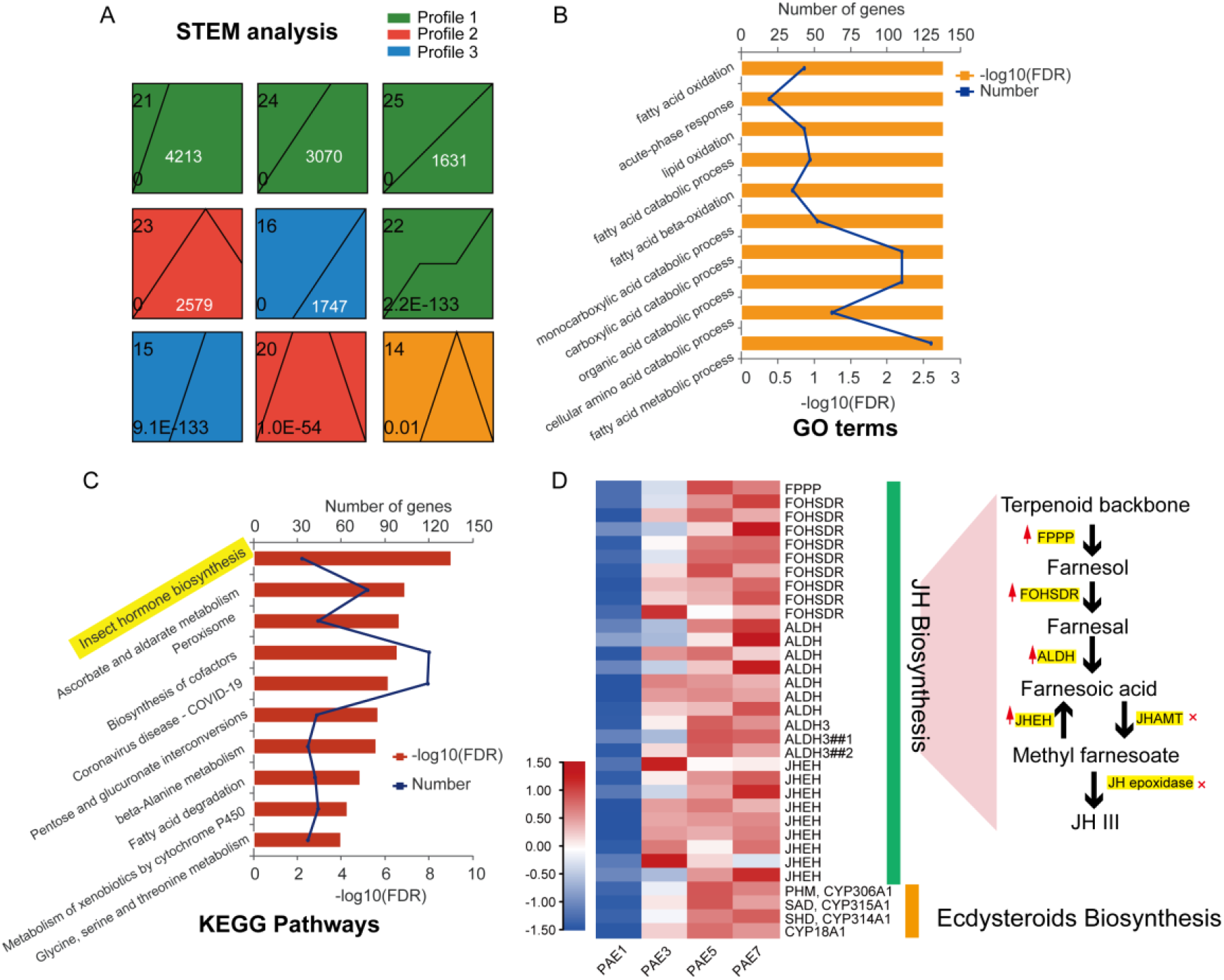
Temporal changes of gene expression based on STEM analysis. (**A**) Nine significant patterns of gene expression in four main possible model profiles recognized by STEM. The patterns (pattern numbers shown at top left) were list from left to right in a descending order of *p* values shown at bottom left in each panel. (**B**) The GO enrichment for genes pooled in patterns 21, 24 and 25. (**C**) The KEGG analysis for genes pooled in pattern 21, 24 and 25. Insect hormone biosynthesis is highlighted in yellow because of its highest –log_10_ (FDR) value. (**D**) Heat map of the expression of the 33 enriched genes in insect hormone biosynthesis pathways in tracheal tubes at four time points. Gene names are shown on the right. The red arrows at the right panel show up-regulation of four enzymes in JH biosynthesis pathway.

### Nematode loading into vector beetle tracheal tubes exponentially increased after third day post eclosion

In order to examine the time and demography of nematodes’ entry into the tracheal system of the vector beetle, we reared beetle pupae on sawyer dust infested with PWN. The number of nematodes in beetle tracheae were counted after the same four time points after adult eclosion (PAE 1, PAE 3, PAE 5 and PAE 7) used in the tracheal transcriptomic analyses (**Figure 5A**). The total number of nematodes in the early stages of tracheal maturation (PAE 1 and PAE 3), were similarly low (less than 400 nematodes per beetle), but increased exponentially after five days post eclosion (PAE 5) (median of 5216 nematodes per beetle, i.e. 15 times higher than after three days post eclosion, and was highest after seven days post eclosion (**Figure 5B**). Therefore, the results indicated that PWN begin to enter the tracheal system three days after the beetles emerged, the loading peak occurring between five to seven days post eclosion during the one-week observation period. Interestingly, the entry of nematodes coincided with the activation of the STEM-enriched insect hormone biosynthesis pathway (Figure **4C** and **4D**), i.e. the enzymes that were mostly up-regulated after three days post eclosion with continuous high expression until the end of the experiment. Such association implies that insect hormones or their precursors may drive nematodes’ entry into tracheal system of vector beetles.

**Figure 5.**
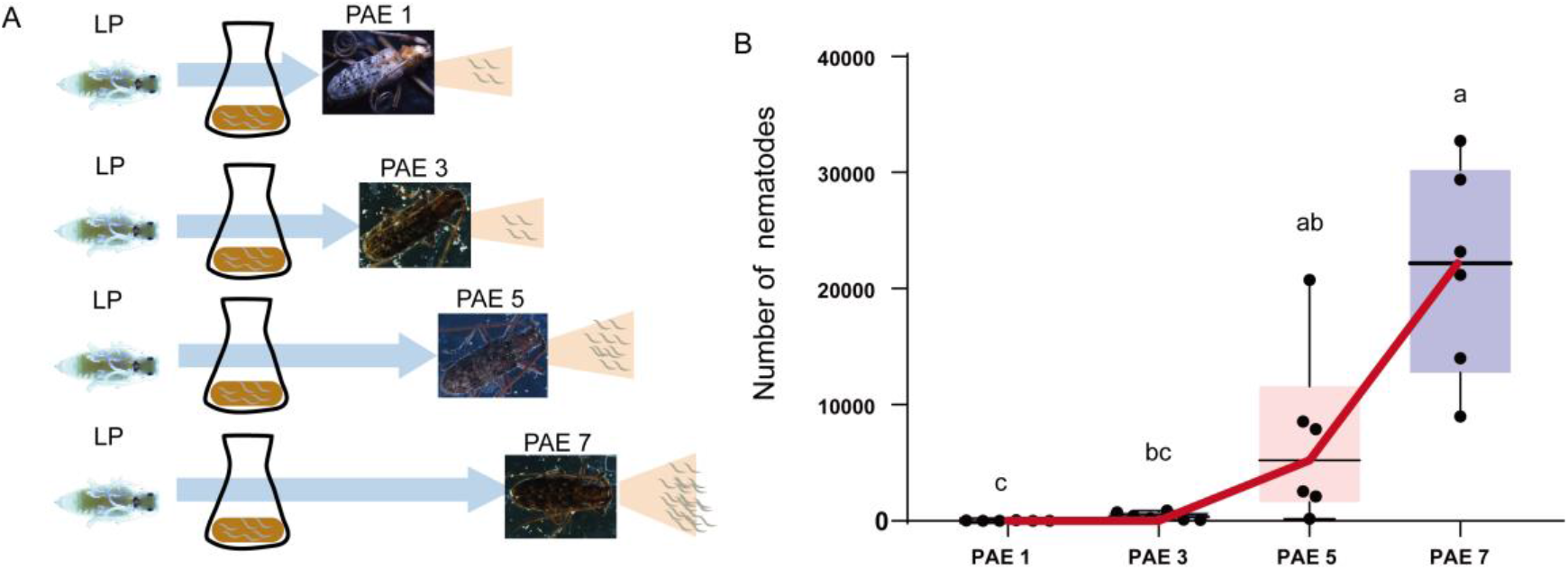
Nematode loading assay. (**A**) Schematic presentation of nematode loading assay, where LP stage beetle pupae (left) were places into flasks containing PWN infested sawyer dust, and kept there until eclosion. The total number of nematodes in the beetle tracheal system were counted from dissected beetles at four different post eclosion time points (day 1= PAE 1, day 3= PAE 3, day 5 = PAE 5 and day 7 = PAE 7) (**B**) Total number of nematodes in tracheal tubes after the four post eclosion time points (*N* = 7 adults for each treatment). The internal lines represent medians, edges of the boxes represent interquartile ranges, and whiskers represent minimum and maximum ranges. Different letters above the boxplot indicate statistically significant differences in the mean relative abundance (Kruskal–Wallis nonparametric test, Dunn’s multiple comparison test, *P* < 0.0001).

### The identification of proteins associated with nematode entry

In addition to small-molecule compounds (i.e. organic fatty acids, or farnesol or farnesal) that serve as potential attractants or stimulants to nematodes, proteins secreted by trachea or residing on tracheal surface might also influence PWN behavior. Therefore, we further identified these proteins based on the DEGs discovered previously. Those genes that were up- or down-regulated during both five and seven days after eclosion (PAE 5 and PAE 7), i.e. during the drastic increase in tracheal PWN loading (**Figure 5 A and B**), were considers as candidate genes for influencing PWNs tracheal attraction and entry. Based on this criterion, 45 up-regulated genes (**Figure 6A**) and 136 down-regulated genes (**Figure 6C**) were identified as potential candidate genes affecting PWN attraction and loading based on their DEG values. For the 45 up-regulated genes, all samples were clearly clustered into two groups based on FPKM profile, high FPKM values found after five and seven days post eclosion (PAE 5 and PAE 7) and lower values found after one and three days post eclosion (PAE 1 and PAE 3) (**Figure 6B)**. Similar clustering pattern was observed for down-regulated genes: tracheal samples collected after five and seven days post eclosion (PAE 5 and PAE 7) had lower FPKM values compared to samples collected after soon after post eclosion (PAE 1 and PAE 3) (**Figure 6D**). Furthermore, we scrutinized the 171 DEGs for tracheal secreted and membrane proteins based on their nr, KOG and pfam annotations, and found that nine secreted proteins and seven membrane proteins were either up- or down-regulated at PAE 5 or PAE 7 (**Figure 6E**, Supplementary table 3). Out of the nine secreted proteins, seven were down-regulated, including two endocuticle structural glycoproteins (SgAbd-2-like and SgAbd-8-like), spidroin-like proteins, angiopoietin-related protein, nidogen, multidrug resistance-associated protein and collagen alpha-1(XV) (**Figure 6E**). Only two of the secreted proteins, defensin-1-like and COL13, were up-regulated after five and seven days post eclosion. Notably, the expression of defensin-1-like had the highest FPKMs (Log_10_FPKM > 3) value among all the 45 up-regulated DEGs, increasing by more than 10 folds at PAE 5 compared with that of PAE 3 (Supplementary table 3). For membrane proteins, the down-regulated proteins included two structural proteins (extensin-2-like and cadherin-89D), and a channel protein (sup-9), whereas the up-regulated membrane proteins included a glutamate receptor (ainate 2-like), two nutrient transporters (Tret1 and NAAT1) and sporozoite surface protein 2-like. Therefore, it is likely that all these 16 proteins which expression co-varied with the number of nematodes entering the tracheal system, influence the interaction between beetle tracheae and PWN during nematode loading.

**Figure 6.**
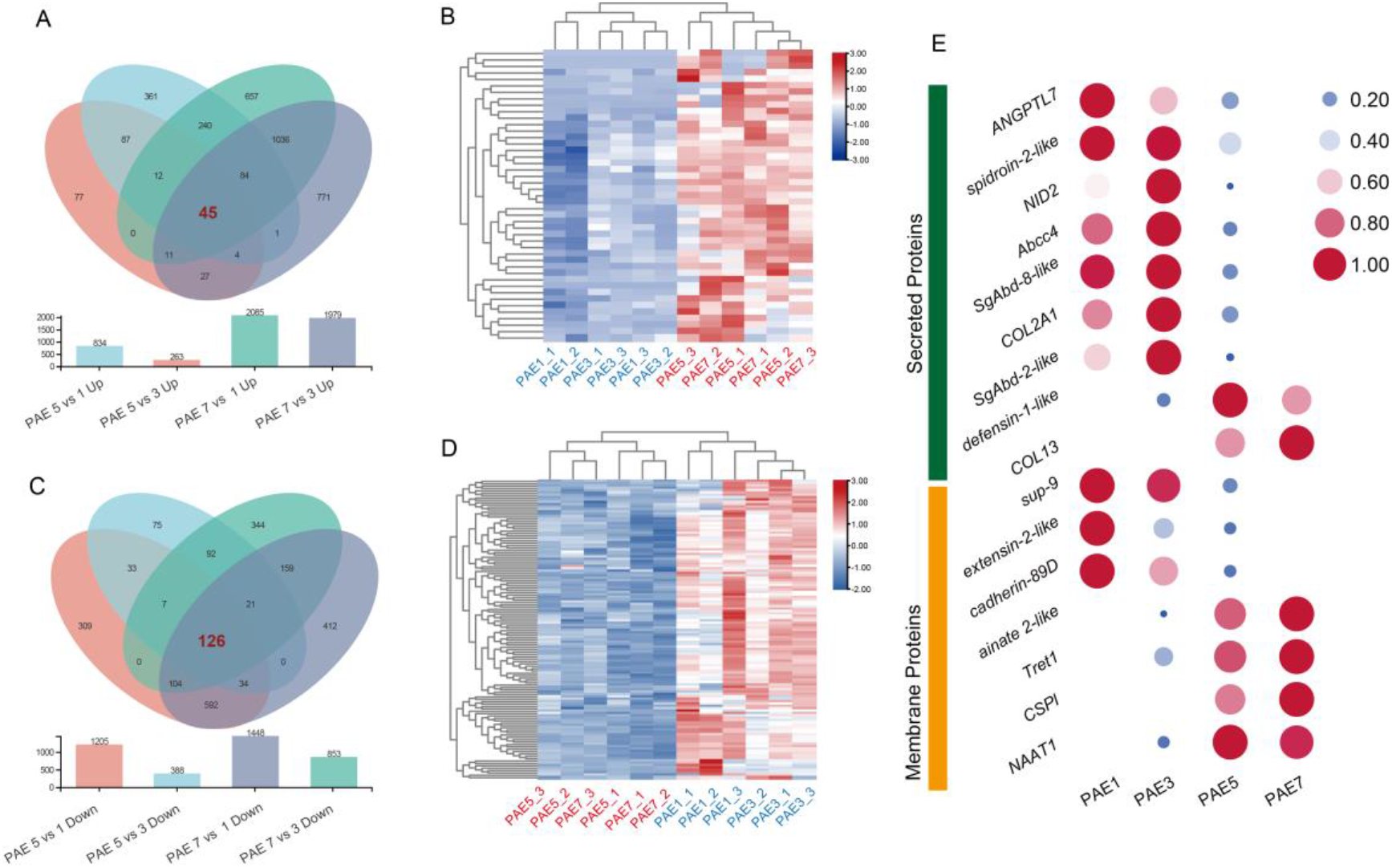
Tracheal candidate proteins for mediating nematode loading. (**A**) Venn diagram of DEGs for up-regulated genes in pairwise comparisons among different post eclosion time points (one day = PAE 1, three days = PAE 3, five days =PAE 5, seven days = PAE 7). The DEGs up-regulated at PAE 5 and PAE 7 are highlighted with red font. The number of DEGs in pairwise comparison between the different post eclosion time points are shown in the bar plot below. (**B**) Heat map of the 45 genes up-regulated at PAE 5 and PAE 7. (**C**) Venn diagram of DEGs for down-regulated genes in pairwise comparisons among different eclosion time points. The DEGs down-regulated at PAE 5 and PAE 7 are highlighted with red font. The number of DEGs in pairwise comparison are shown in the bar plot below. (**D**) Heat map of the 126 genes down-regulated at PAE 5 and PAE 7. (**E**) Heat map showing tracheal secreted and surface proteins specifically up- or down-regulated at PAE 5 and PAE 7. Both the size and the color of the dots indicate the FPKM value, larger, dark red spots indicating strong up-regulation and small bark blue spots indicating strong down-regulation. Protein names are shown on the left side.

## Discussion

Our results demonstrate exquisite synchronization of tracheal entry of a plant parasitic nematode with tracheal development and maturation of its vector beetle. We found that newly emerged adult beetles experience programmed tracheal growth at both morphological and gene expression level, which is followed by activation of insect hormones biosynthesis. The rapid increase in the number of PWNs in vector’s trachea after five days post eclosion indicates that the formation of primary tracheal branches is a prerequisite for PWN entry, and that hormone-derived compounds from beetle tracheae during this period may serve as attractants to PWN. Tracheal up-regulation of Defensin-1-like and down-regulation of some secreted proteins after five and seven days post eclosion may further facilitate the entry of PWN. This study reveals the reliance of parasites on the timing of tissue development for their successful entry into their hosts or vectors, and suggest possible molecular mechanisms underlying PWN entry into tracheal systems of *Monochamus* spp. vector beetles.

This study assessed for the first time the morphological and genetics basis underlying tracheal growth after eclosion in a holometabolous insect, and its role for the entry of plant-parasitic nematode the insect vectors. Interestingly, we found that despite the most dramatic changes in tracheal gene expression of *M. alternatus* occurred after five days post eclosion, the tracheal genes were expressed sequentially in modules throughout the sclerotization period to regulate the maturation of tracheal systems, a pattern which is similar to the tracheal development demonstrated on *D. melanogaster* embryos (Hayashi and Kondo, 2018). More specifically, we found that Trh, Vvl and Bnl-Btl signaling were activated during the first day post eclosion. Both Trh and Vvl are known to be the earliest upstream regulatory transcription factors activating numerous tracheal genes in insects (Hanna and Popadic, 2020; Hayashi and Kondo, 2018), whereas Bnl-Btl signaling is known to be important for guiding tracheal branch migration, as well as being involved in the establishment of tracheal pattern at multiple developmental stages (Hayashi and Kondo, 2018). In addition, transcriptional factors involved in shaping the tracheal tubes, such as *knirl* and *salm*, were also up-regulated soon after eclosion, collectively indicating that vigorous formation of primary branches occurred during the first day after the adult beetle eclosed. Interestingly, these genes that were up-regulated soon after eclosion were rapidly down-regulated during three or five days after eclosion. Instead, genes associated with terminal branching and branch fusion were up-regulated during this period, demonstrating a trade-off between the expression genes governing primary and terminal branching. During the first post eclosion day, we also observed up-regulation of two key chitin-modifying enzymes (Vermiform and Serpentine) that drive the assembly of a luminal matrix, and found an increase in tracheal tubular diameter between the first and third day post eclosion. Therefore, our results suggest that the tracheal diametric expansion occurs almost synchronically with primary branching right after adult eclosion. This differs from the patter observed in *Drosophila* embryos, in which this process seem to be sequential occurring within a defined time interval, starting with primary branching, later followed by embryonic tube dilation (Zuo et al., 2013). Moreover, contrasting with the continuous tube elongation occurring during insect embryogenesis (Hayashi and Kondo, 2018), genes regulating tracheal tube length after adult eclosion (such as *yokie, aPKC, rhoA, shrb, yrt* and *scrib*) were up-regulated during third or fifth day post eclosion with continuous high expression until the end of the experiment. These differences of tracheal gene expression imply modification of gene regulation during metamorphosis compared to embryogenesis, perhaps due to the need to rapidly form a complex tracheal network that the mobile adult insect needs for sufficient oxygen transportation (e.g. during flight).

Our results also showed that the tracheal entry of PWNs is reliant on the developmental stage of the beetle’s tracheal systems. Before the third day after eclosion, only a few PWNs entered the trachea, despite that the newly enclosed adults had well-developed external adult morphology. The abstention of PWNs to enter the tracheae at this stage is likely due to the tracheal immaturation preventing efficient nematode loading, as the tracheal systems at this stage was likely still undergoing primary branching and diametric expansion. However, after the third day of post eclosion, nematode loading drastically increased, suggesting that the nematode entry require well-constructed primary branches with a large enough diameter, while being less dependent on the tracheal terminal branching and branch fusion. Many parasites show tropism towards certain host tissues (Silva Pereira et al., 2019), and our study provided empirical evidence that tissue maturation is likely a key factor affecting parasite entry, explaining the temporal fidelity of PWN to enter vector trachea.

Trachea may also participate in the insect’s hormone regulation by producing and/or storing pheromones needed in the tracheae or in other tissues. A previous study has shown that in *Drosophila spp*., corpora allata that produces JH, prothoracic glands that produce ecdysteroids, and trachea, share a common developmental origin (Sanchez-Higueras et al., 2014). In our study, we found that both JH and ecdysteroids biosynthesis pathways which are involved in insect hormone biosynthesis, were significantly enriched during tracheal maturation after the first post eclosion day based on the STEM analysis. Thirty-three key enzymes involved in JH and ecdysteroids biosynthesis were the significantly up-regulated during the tracheal post eclosion maturation, indicating that trachea may serve as a reservoir of JH and ecdysteroids or their precursors, thus being functionally analogous to endocrine glands. The biological significance of these trachea-originated hormones or their precusors, however, was not tested in this study, warranting further investigation. Because ecdysone signaling is required in the formation of dorsal branches in *Drsophila* embryos (Taira et al., 2021), and based on the enrichment of these genes found also in this study, it is also likely that these hormones also are required in post eclosion tracheal maturation in *Monochamus spp*. beetles. Alternatively, it is possible that besides corpora allata and prothoracic glands, beetle trachea may serve as an additional supplier of JH and ecdysteroids to regulate biological processes in other tissues, because insect hormones, especially JH, has multiple functions in adult insects. For example, despite its original role as a status-quo hormone regulating insect metamorphosis, JH is also suggested to function as a gonadotropin with pleiotropic effects on diapause and immunity (Li et al., 2019). Furthermore, high hemolymphic JH concentration promotes oogenesis of insect females by inducing vitellogenin synthesis in the fat body and vitellogenin uptake in the follicle cells (Santos et al., 2019), whereas in males, JH participate in courtship behaviour by modulating the sensitivity of olfactory receptor neurons (Lin et al., 2016). Future studies should investigate the temporal and spatial pattern of the expression of downstream genes (e.g. *krh1* and *met*) in adult beetles to test the role of trachea as a JH and ecdysteroid target and/or supplier.

The STEM-enriched genes in the hormone synthesis were significantly up-regulated after third day of post eclosion, which was also the time frame when increase in nematode loading was detected. Such close association of the timing of these events tentatively suggests that vector tracheal hormones or their precursors may act as kairomones that mediate the behavior or physiology of PWN. On one hand, due to lack of JHAMT in beetle trachea that generally participates in the production of JH III, terpenoids precursors including farnesol, farnesal and farnesoic acid are produced in JH biosynthesis. These compounds can likely functions as attractants to PWN, because several terpenoids, including pinene and longifolene released by beetle larvae and pine trees, are known to be attractive to PWN (Zhao et al., 2007). On the other hand, JH precursors and ecdysteroids released by the beetle trachea may induce a priming effect that elicit an array of physiological responses in PWN. For example, farnesol and ecdysone have been shown to have stimulatory effect of on various species of nematodes (Barker and Rees, 1990; Lawrence, 1991), including farnesol mimicking the effect of juvenile hormone and stimulated DNA and RNA synthesis in some parasitic nematodes. In addition, exposure to ecdysone and 20-E have been shown to promote moulting and stimulating meiotic reinitiation in oocytes and microfilarial production in filariae in various nematode species (Barker and Rees, 1990; Lawrence, 1991).

In addition to potential kairomones that mediate nematodes’ behavior, tracheal secreted or membrane proteins may also influence the behavior of PWN after the nematode has entered the tracheal system. Defensin and Defensin-like antimicrobial peptides (AMPs) are well known as innate immune factors against bacterial infections in animal respiratory systems (Travis et al., 2001). Up-regulation of *defensin-1-like* in trachea after five and seven days post eclosion therefore suggests a presence of defense against air-borne bacterium during the later stages of tracheal development. Recently, Defensin has been shown to assists the colonization of an entomopathogenic fungus on insect body surface by manipulating bacterial load on the insect surface (Hong et al., 2022). Our current study demonstrated a positive correlation between Defensin expressional level and PWN number in the tracheae. In addition, our previous results found a significantly higher *defensin-1-like* expression in tracheae loaded with PWN compared to if infected with an entomopathogenic nematode (Zhou et al., 2018). Therefore, Defensin-1-like possibly promotes the entry of PWN by regulating bacterial community in tracheal tubes. Besides the beneficial components that promote PWN entry during the later stage of post eclosion (PAE 5 and PAE 7), some proteins (such as glycoproteins, collagen) were down-regulated. Because these secreted proteins are known to form physical barriers against pathogens (Tomlin and Piccinini, 2018), down-regulation of these components may also mediate PWN entry by preventing their potential negative effects on the attachment of PWN.

In summary, our study provides in-depth insights into the behavioral adaptation of vector-transmitted parasites in response to target tissue development of their vectors, and offers potential molecular targets that would assist the control of pine wilt disease.

## Materials and methods

### Nematodes and beetles

Mature larvae of *M. alternatus* were acquired from dead pine trees in the Fuyang area, Hangzhou City, Zhejiang Province, and reared in an incubator (26°C, humidity 35%) until pupation. *B*.*xylophilus* were originally collected from infected trees in Shaanxi Province and reared with *Botrytis cinerea* in potato dextrose agar (BD, 213400) plates at 25°C for generations until used in the present study.

### Measurement of tubular size

Adult beetles post one, three, five and seven days after eclosion were dissected by ventral filleting. The morphological traits of thoracic tracheal tubes were observed under a stereomicroscope (Olympus SZX16, Japan) and were imaged by microscope digital camera software (DP2-BSW), then the diameters of tracheal tubes were measured by the software (DP2-BSW).

### Nematode loading assay

*B. cinerea* kept on autoclaved (121°C, 30min) barley medium (10g barley in 15ml water) in 50ml Erlenmeyer flasks, were inoculated with 1,000 propagative nematodes. Five days after the nematode inoculation, pine wood sawdust layer was added on top of the barley layer and kept for another two weeks. Beetles were placed on the nematode-infected sawdust layer at late pupa stage until dissection after eclosion. Emerged adult beetles were dissected by ventral filleting, and the tracheal tubes located on both sides of the thorax were dissected under a stereomicroscope (Olympus SZX16, Japan). The excised tracheal tubes and the left part of the body were washed in PBST (Solarbio, P1033) to irrigate the nematodes and presence of nematode was confirmed under a stereomicroscope (Olympus SZX16, Japan). The total number of isolated nematodes per beetle were counted in plates with an inverted microscope (Olympus CKX41, Japan).

### RNA extraction

Upon pupation, twelve beetles were assigned randomly to four groups. They were checked and recorded for their eclosion status on daily basis. The four groups were sampled at day 1, day 3, day 5 and day 7 respectively post eclosion. The thoracic tracheal trunks were dissected carefully, washed with PBS to get rid of nematodes, and immediately flash frozen with Trizol (Invitrogen,15596018) and stored in -80°C before RNA extraction.

Total RNA was isolated from the frozen samples using Trizol reagent and genomic DNA was removed using DNase I (TaKara). The integrity and purity of the total RNA quality was determined by 2100 Bioanalyser (Agilent Technologies) and quantified using the ND-2000 (NanoDrop Technologies). Only high-quality RNA sample (OD260/280=1.8∼2.2, OD260/230≥2.0, RIN≥8.0, 28S:18S≥1.0, >1μg) was used to construct sequencing library.

### Library preparation and sequencing

RNA purification, reverse transcription, library construction and sequencing were performed at Shanghai Majorbio Bio-pharm Biotechnology Co., Ltd. (Shanghai, China) according to the manufacturer’s instructions (Illumina, San Diego, CA). The ranscriptome library was prepared following TruSeqTM RNA sample preparation Kit from Illumina (San Diego, CA) using 1μg of total RNA.

### De novo assembly and annotation

The raw paired end reads were trimmed and quality controlled by fastp (https://github.com/OpenGene/fastp) with default parameters. Then clean data from the samples were used to do de-novo assembly with Trinity (http://trinityrnaseq.sourceforge.net/). Then the assembled transcripts were assessed and optimized with BUSCO (Benchmarking Universal Single-Copy Orthologs, http://busco.ezlab.org), TransRate (http://hibberdlab.com/transrate/) and CD-HIT (http://weizhongli-lab.org/cd-hit/). All the assembled transcripts were searched against the NCBI protein non-redundant (NR, ftp://ftp.ncbi.nlm.nih.gov/blast/db/), Swiss-Prot (http://web.expasy.org/docs/swiss-prot_guideline.html), Pfam (http://pfam.xfam.org/), Clusters of Orthologous Groups of proteins (COG, http://www.ncbi.nlm.nih.gov/COG/), GO (http://www.geneontology.org) and KEGG (Kyoto Encyclopedia of Genes and Genomes, http://www.genome.jp/kegg/) databases using BLASTX to identify the proteins that had the highest sequence similarity with the given transcripts to retrieve their function annotations and a typical cut-off E-values less than 1.0×10-5 was set.

### Differential expression analysis and functional enrichment

To identify DEGs (differential expression genes) between two different samples/groups, the expression level of each gene was calculated according to the fragments per kilobase of exon model per million mapped fragments (FPKM) method. RSEM (http://deweylab.biostat.wisc.edu/rsem/) was used to quantify gene abundances. Essentially, differential expression analysis was performed using the DESeq2, DEGs with |log_2_(foldchange)| ≥ 1 and *P*-adjust ≤ 0.05 were considered to be significantly different expressed genes. In addition, functional-enrichment analysis including GO (Gene Ontology, http://www.geneontology.org) and KEGG (Kyoto Encyclopedia of Genes and Genomes, http://www.genome.jp/kegg/) were performed to identify which DEGs were significantly enriched in GO terms and metabolic pathways at P-adjust ≤ 0.05 compared with the whole-transcriptome background. GO functional enrichment and KEGG pathway analysis were carried out by Goatools (https://github.com/tanghaibao/Goatools) and KOBAS (http://kobas.cbi.pku.edu.cn/ home.do)

### Identification of tracheal genes, secreted and membrane proteins

For tracheal genes, the amino acid sequences of known tracheal genes from *Drosophila melanogaster*, retrieved from flybase were used as inquries for a Blastn search of *M. alternatus* predicted protein. We manully confimed the potential candidates of *M. alternatus* tracheal genes based sequence similarity with *Drosophila* homologues and nr, pfam or KOG annotation. To identify the secreted and membrane proteins, we scrutinized gene sets for nr, pfam or KOG annotation. Their location was determined based on localization information in Uniprot database (https://www.uniprot.org/).

### Statistical analysis

The sample size is determined according to previous publications in the pine sawyer beetle species (Wu et al., 2019; Zhao et al., 2016; Zhou et al., 2018). For nematode loading, individuals were randomly allocated into experimental group and control group, and no restricted randomization was applied. All experiments were performed for at least 3 independent biological replicates.

Gene expression patterm analysis was perform by Short Time-series Expression Miner software (STEM) (Ernst and Bar-Joseph, 2006) on the Majorbio Cloud Platform, a free online platform for data analysis(www.omicshare.com/tools). The Maximum Unit Change in model profiles between time points is 1. Maximum outmput profiles number is 50. Minimum ratio of fold change of DEGs is no less than 2.0.

A Kolmogorov Smirnov test was used to check the normality of data. One-way ANOVA with Dunn’s or Tukey’s multiple comparison were used for comparisons of tubular diameter. Kruskal–Wallis with Dunn’s multiple comparison were used for comparisons of nematode loading number. Statistical analysis was performed with Graphpad Prism 8.0 (Graphpad software, San Diego, California, USA).

## Acknowledgements

This work was funded by the National Natural Science Foundation of China (32088102, 32061123002) and National Key Research and Development Program of China (2021YFC2600100).

## Author contribution

X.T. and J.S. conceived and designed the experiments. J.S. supervised the study. X.T. performed loading and counting of PWN, tracheal dissection, tracheal observation and transcriptomic analysis. X.T. prepared the manuscript draft. X.T., T-M. K. and J.S. revised the manuscript.

## Additional files

**Supplementary table 1** Gene list of 55 genes related to tracheal development in *M. alternatus*.

**Supplementary table 2** Gene list of 33 genes enriched in insect hormone biosynthesis pathway were related to biosynthesis of Juvenile hormone (JH) and ecdysteroids.

**Supplementary table 3** Gene list of tracheal secreted and membrane proteins specifically up- or down-regulated at PAE 5 and PAE 7.

## Data availability

All data generated or analyzed during this study are included in the manuscript and supporting file, The following dataset was generated:

| Author(s) | Year | Dataset title | Dataset URL | Database and Identifier |
| --- | --- | --- | --- | --- |
| Tang X | 2022 | RNA-Seq of tracheal tubes in adult pine sawyer beetle | https://ngdc.cncb.ac.cn/gsa/s/fEsL0f8r | bigd, CRA008617 |

